# Asymmetric Reconstruction of the Aquareovirus Core at Near-Atomic Resolution and Mechanism of Transcription Initiation

**DOI:** 10.1101/2022.08.31.505870

**Authors:** Alexander Stevens, Yanxiang Cui, Sakar Shivakoti, Z. Hong Zhou

## Abstract

The *Reoviridae* family of dsRNA viruses is characterized by its members’ capacity for endogenous transcription of their multipartite genomes within proteinaceous capsids of 1 to 3 layers. These viruses share inner core particles (ICPs) that conform to icosahedral, T=2*, symmetry, but differ in two major respects: first, the presence or absence of RNA-capping turrets at each icosahedral vertex; second, the number of additional host-specific capsid layers that are often lost upon cell entry. While the role of these additional layers in host infection is generally understood, the absence of asymmetric ICP structures from turreted, multilayered reoviruses has obfuscated our understanding of how successive removal of these external layers impact the structural organization of the ICP and transcription initiation. Here, we present the 3.3 Å resolution structure of the aquareovirus (ARV) ICP, and atomic models of the capsid proteins VP3 and VP6, transcriptional enzymatic complex (TEC) subunits VP2 and VP4, and RNA-capping turret protein VP1. These structures reveal significant differences when compared to those of the coated ARV, as well as their counterparts in single-layered cytoplasmic polyhedrosis virus (CPV). Compared to the double-layered ARV virion and infectious subvirion particle structures, the ARV ICP undergoes significant capsid expansion and widening of the nucleotide processing channels in its TEC and turret. Thus, the loss of outer capsid layers may regulate transcription initiation in ARV, unlike CPV which relies solely on allosteric regulation by binding transcriptional cofactors. These results shed new light on the mechanism of transcription initiation amongst turreted, multilayered members of *Reoviridae*.

## Introduction

Aquareovirus (ARV) is one of 15 genera from the *Reoviridae* family of double-stranded RNA (dsRNA) viruses (collectively called reoviruses), which causes hemorrhagic disease in economically important aquaculture of North America and Asia. *Reoviridae* members infect a wide range hosts and include the human borne rotavirus which causes approximately 128,000 deaths in young children each year (Troeger et al., 2018). As reoviruses are non-enveloped, host-cell susceptibility is determined by their proteinaceous capsids which enclose multisegmented genomes and are most readily distinguished by the number of icosahedral capsid layers, and the presence (*Spinareovirinae* subfamily, 9 genera) or absence (*Sedoreovirinae* subfamily, 6 genera) of mRNA capping turrets along the innermost layer (King et al., 2012). Member viruses of both *Sedo-* and *Spinareovirinae* enclose their segmented dsRNA genomes within capsids of one to three concentric layers but, regardless of the number of genome segments, capsid layers, or turrets, their innermost layer from is an icosahedral, T=2*, inner capsid particle (ICP). During cell entry, reoviruses often shed their outer layers leaving the ICP intact to evade the host’s viral genome detection machinery, and act as nanomachines to endogenously transcribe positive-sense viral messenger RNA (mRNA) at transcriptional enzymatic complexes (TECs) situated underneath the icosahedral vertices within the ICP (Cui et al., 2019; Ebert et al., 2002; Skehel and Joklik, 1969).

Among turreted reoviruses, CPV is the simplest, structurally, possessing a single-layered capsid equivalent to the ICP of double-or triple-layered reoviruses but which is also the intact virion (Hill et al., 1999; Zhang et al., 1999; Zhou et al., 2003). Owing to its simple structural organization and more complete characterization, CPV has long served as a model to study the lifecycles of other turreted reoviruses. When presented a susceptible cell, a quiescent CPV virion enters via endocytosis (Chen et al., 2018). Upon cell entry, the CPV turret proteins (TPs) bind S-adenosyl methionine (SAM) and adenosine triphosphate (ATP), inducing subtle capsid expansion along with architectural changes to the TECs that facilitate transcription (Yang et al., 2012; Yu et al., 2015; Zhang et al., 2022b). In the transcribing state, viral RNA-dependent RNA polymerases (RdRps), along with nucleotide triphosphatases (NTPases), transcribe viral mRNA, which receives its 5’ cap from the turrets en route to the cytosol where they utilize the host’s translational machinery.

However, the single-layered architecture that makes CPV an attractive model system also renders it an inadequate model to describe the numerous, multi-layered, *Spinareovirinae* members such as the double-layered aquareoviruses (ARVs) whose outer layer(s) may prevent the same conformational changes to the inner capsid (Yang *et al*., 2012; Yu *et al*., 2015; Zhang *et al*., 2022b). Endemic aquareoviruses in North America and China cause hemorrhagic diseases in golden shiner and grass carp, respectively (Fang et al., 2005; Nason et al., 2000); and the cryo-electron microscopy (cryoEM) structures of the American and Chinse aquareovirus virion isolates have been resolved to 22 and 3 Å resolution, respectively (Ding et al., 2018; Shaw et al., 1996; Wang et al., 2018). These double-layered structures of ARV are highly similar to those of the mammalian orthoreovirus (MRV) virion (Dryden et al., 1993; Pan et al., 2021), but are distinct to those of CPV. The second layer of ARV capsids give them an additional life stage between the quiescent and transcribing stages known as the primed state, which involves a more complicated array of structural changes and outer layer removal to prepare the virus for transcription (Ding *et al*., 2018).

The outer ARV capsid layer is comprised of small VP7 protection proteins and large VP5 penetration proteins. During infection, VP7 is cleaved extracellularly (Zhang et al., 2022a), exposing VP5 and creating a maximally infectious form of ARV known as the infectious subvirion particle (ISVP) (Nason et al., 2000). The ISVP uses the newly exposed penetration proteins to escape the endocytic pathway and deposit the transcriptionally competent ICP to the host cytosol (Fig. S1) (Kim et al., 2004; Nason *et al*., 2000; Skehel and Joklik, 1969; Zhang et al., 2010). Within the cytosol, the ICP can simultaneously synthesize and export viral mRNA from all genome segments and append 5’ methylated guanosine caps to the nascent mRNA. This T=2* icosahedral core is composed of 120 copies each of the capsid shell protein (CSP) VP3 and the clamp proteins VP6. Each ICP encloses the 11 dsRNA segments and their accompanying transcriptional enzymatic complexes (TECs), each which is a heterodimer of RdRp VP2 and its cofactor, NTPase VP4. Asymmetric sub-particle reconstruction methods coupled with iterative symmetric relaxation workflows have enabled visualization of the asymmetrically attached TECs within the core of several dsRNA viruses (Cui *et al*., 2019; Ding *et al*., 2018; Zhang et al., 2015). To date, only the asymmetric structures of complete ARV virions and ISVPs have been resolved to high-resolution, thus it is unclear what, if any, structural changes occur within the ICP during uncoating (Ding *et al*., 2018; Wang *et al*., 2018). Consequently, whether the external layers of the multi-layered turreted dsRNA viruses also play a structural and/or functional role in transcription regulation in addition to their involvement in cell entry is not known in the absence of a structure for comparison of ICPs to virions.

In this study, we report the first asymmetric reconstruction of the ARV ICP by cryoEM. Comparison with existing ARV virion and ISVP structures reveals subtle architectural changes between the coated and uncoated core particles which may act to facilitate endogenous transcription. These changes yield a particle with significantly increased internal volume and widened mRNA exit channels. Our structure also enabled modeling of previously unresolved TEC and CSP segments, which illustrate the interactions between the CSP N-termini, the TEC, and the adjacent genomic RNA absent the protective VP7-VP5 layer. As the first high-resolution asymmetric structure of the innermost core of a multi-layered member of the *Spinareovirinae* subfamily of *Reoviridae*, this structure fills in a knowledge gap in the ever-growing repository of asymmetrically reconstructed dsRNA viruses, demonstrating the conformational difference between coated and uncoated reoviruses, and informing on how those changes may help to regulate transcription.

## Results and Discussion

### Asymmetric reconstruction of GSRV core resolving 11 TEC-occupying and 1 RNA-filled vertices

We determined the asymmetric structure of the GSRV ICP by following a sequential symmetry relaxation approach as described previously (Ding *et al*., 2018; Zhang *et al*., 2015). Virus structures were initially refined with icosahedral symmetry imposed and finer details of the non-icosahedral related regions were resolved via symmetry expansion and focused classification as detailed in previous works (Ding *et al*., 2018; Pan *et al*., 2021) (Materials and Methods).

Following this procedure, four asymmetric structures were obtained: three from the sorted I5 vertices, which include the polar vertices at 3.3 Å and the tropical vertices with and without the TECs to 3.3 Å and 3.7 Å respectively, along with a reconstruction of the complete capsid featuring 11 TECs at 4.2 Å. The resolution of TEC containing sub-particles following averaging of equivalent copies reached 3.4 Å. Our asymmetric reconstruction of the GSRV core contains 60 VP1 turret proteins, 120 VP3 capsid shell proteins (CSPs), 120 VP6 clamp proteins, 11 TECs under 12 vertices, with only one southern tropical vertex lacking a TEC complex (Fig. 1F).

**Figure 1.**
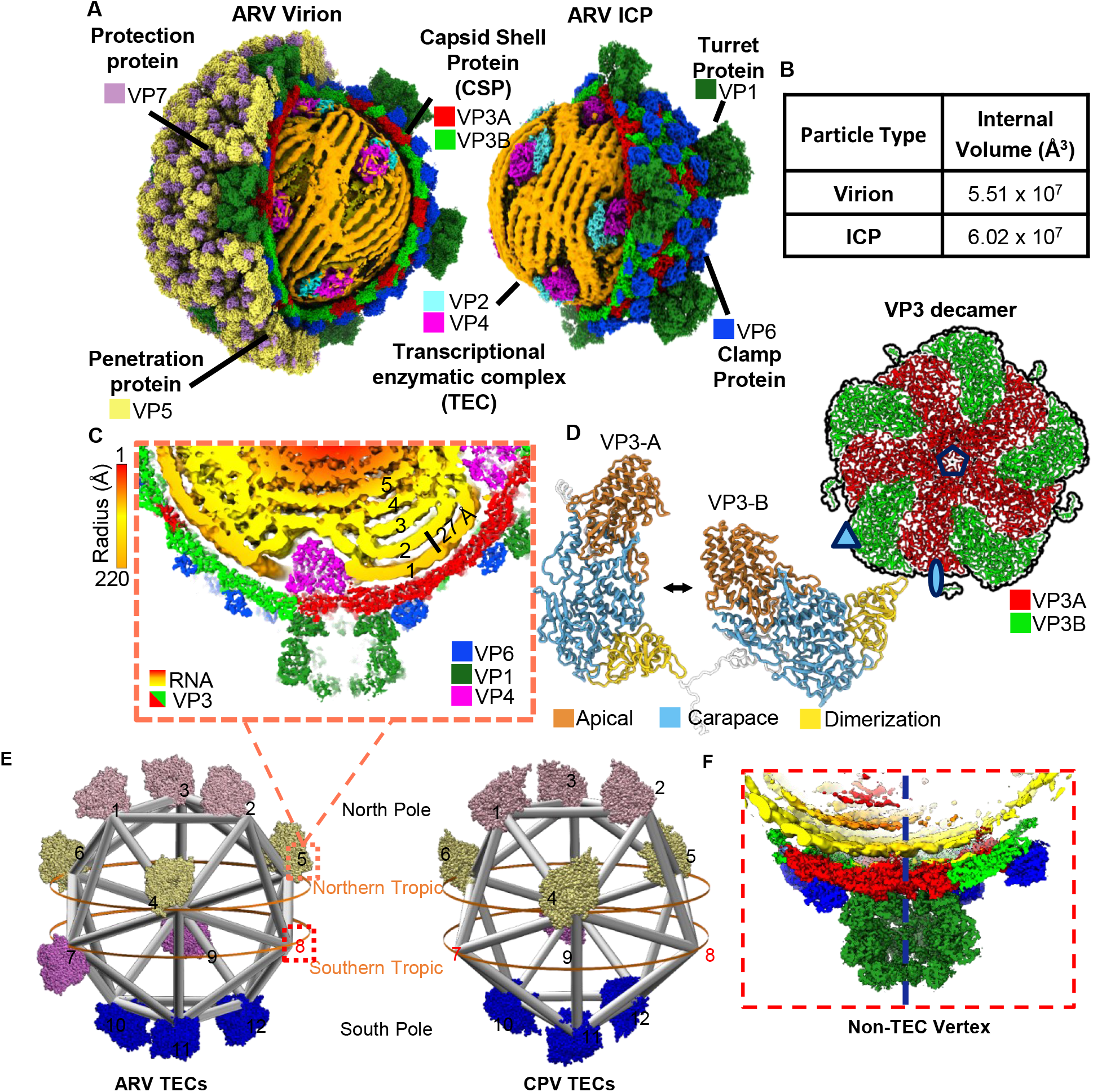
Asymmetric reconstruction of ARV ICP. (A) CryoEM structures of ARV virion (left) (5VST) and ICP (right) with portion of their external layers removed along hemispheres to expose genome and TECs. (B) Table comparing volumetric difference between virion and ICP lumen. (C) Cross sectional view of Tropical Vertex featuring TEC and a turret. Layers of genome segments are numbered from 1-5 based on proximity to capsid wall. (D) Ribbon Diagram of ICP capsid decamer (top right) and VP3_A_ (left) and VP3_B_ (right) conformers with domains indicated and colored differently. (E) Position and orientation of the 11 ICP TECs relative to an icosahedron (grey) with Northern and Southern “poles” and “tropics” indicated. For clarity, RdRp distribution model not drawn to scale. (F) Cross sectional view of ICP vertex without TEC showing that the TEC-free vertex is occupied by RNA density (yellow).

These 11 TECs within GSRV core are internally organized following pseudo-*D*_*3d*_ symmetry, with 6 TECs occupying the icosahedron’s poles while 5 are situated along the equatorial plane at the northern and southern tropics (Fig. 1D). 11 dsRNA segments between 0.8 and 3.9 kbps compose the GSRV genome and are tightly packaged inside the capsid (Fig. 1C); the outermost dsRNA layer shows large, persistent, parallel lengths for each RNA double-strand, with an average of 27 Å between neighboring strands, consistent with the previous ARV structures, despite the increased internal capsid volume (Fig. 1B-C) (Ding *et al*., 2018).

### Loss of outer coat proteins leads to expansion of ICP capsid

Lacking the outer capsid proteins (OCPs) VP5 and VP7, the ARV ICP retains the icosahedral, T=2*, inner capsid shell composed of 60 asymmetric dimers of the 1,214-residue, wedge-shaped CSPs and 120 symmetrically arranged copies of the clamp protein (VP6) which form the ICP frame and provide support respectively (Fig. 1A). Consistent with the previously resolved ARV capsid structures, VP3 dimers (VP3_A_ and VP3_B_) encircle each 5-fold vertex; with VP3_A_ conformers seated around the 5-fold vertex center, creating pores adjacent to each TEC’s template exit channel for direct transcript capping and release via VP1 turret proteins (TPs) (Fig. 1C-D). VP3_B_ conformers partially intercalate between VP3_A_ monomers, and form 3-fold vertices with neighboring decameric assemblies (Fig. 1D). Absent conspicuous CSP rearrangements, the internal volume of the ICP increased by about 9.3% relative to the previously resolved GCRV virion and ISVP with which GSRV shares 96-100% a.a. sequence identity (Fig. 1B) (Ding *et al*., 2018; McEntire et al., 2003; Wang *et al*., 2018). The motion undergone by individual decamers includes a 10 Å rise away from the virion origin and subtle expansion of the CSPs (Movie S2) (Goddard et al., 2018). The observed expansion can be attributed to a non-uniform elongation of the CSP monomers approximately 6 Å radially from the I5 vertices, relative to their coated counterparts, and this creates an expanding and rising motion around the decameric subunit (Movie S1,3). Motion isn’t uniform across the CSP monomers however and differs between the VP3 conformers.

Per historic naming conventions, each ARV CSP is divided into three distinct domains. Extending radially from the I5 vertices, these domains include the α-helix rich apical “tip” nearest the I5 vertices (a.a 486-830), the large carapace domain (a.a. 190-485, 831-976, and 1144-1214), and the small β-sheet rich dimerization domains (a.a. 977-1143) (Fig 1D). The local motion of CSP dimers (Movie S2) reveal a non-uniform elongation of the apical and carapace domains, with the alpha helices migrating away from the I5 vertices and towards the icosahedral two-fold and three-fold (I2 and I3) vertices for VP3_A_ and VP3_B_ respectively (Movie S2). In the apical domain helices 13 and 14 and their conjoining loop (a.a. 490-518), which line the I5 transcript exit channel at the luminal side of the capsid, are largely unperturbed by the capsid shifts and help to maintain a consistent pore diameter. By contrast, the other I5 adjacent elements of the apical domains move away from the I5 channel, and the helices 13 and 14 of VP3_**B**_ migrate in a similar manner as the other secondary structural elements. Closer inspection reveals striking conservation of VP3_A_ residues adjacent to the I5 pore (a.a. 490-518, RMSD .464 Å) when compared to the quiescent virion and ISVP structures (Ding *et al*., 2018; Wang *et al*., 2018). Despite exhibiting outward motion, the secondary structural elements of CSPs which interface with the clamp and turret proteins interfaces appear to move as rigid bodies, presumably constrained by interactions with the essential clamp and turret proteins (Movie. S2). The expansion of the CSPs leads to a significant increase of internal capsid volume, from 5.51 ×10^7^ Å^3^ to 6.02 ×10^7^ Å^3^, or about 9% from the ARV virion to the ICP (Fig. 1B). By reducing the packaging density and therefore the viscosity of the genome, the enlarged ICP provides the ridged dsRNA segments greater freedom of motion and presumably lowers the energy required to initiate transcription (Demidenko and Nibert, 2009).

The inner-outer capsid protein (VP3-VP5) interactions of ARV are mediated through the clamp protein VP6, making it important not only for stabilization against the genome but in secondary layer association. Previous work has shown the closely related MRV ICPs can be recoated to form ISVPs (Farsetta et al., 2000), suggesting the ARV ICP clamps remain in a VP5 receptive state following uncoating. Indeed, superimposition of the ICP and virion clamp proteins reveal significant conformational similarity (RMSD 0.667 Å) despite the migration of the clamps away from the I5 vertices in the ICP particles. The architectural changes of VP6 can, therefore, be summarized as shifts of 4-5 Å away from the I5 vertices along the capsid wall, with no conspicuous, irreversible changes that would inhibit OCP VP5 association. So, in addition to its role stabilizing the VP3 ICP against the force exerted by the genome, VP6 provides a platform onto which the VP5 layer can bind and possibly compress core particles to halt transcription. While this dynamic may be inconsequential for parental ARV particles, whose VP5 proteins are irreversibly cleaved during entry, it may be a key mechanism to halt transcription in newly assembled ARV particles following complimentary strand synthesis, as observed in nascent rotavirus particles (Trask et al., 2012).

### Capsid expansion leads to subtle TEC rearrangements

Several dsRNA segments interact with each TEC and these interactions stabilize the genome, enabling improved visualization of their major and minor grooves within the ARV ICP as observed in other RNA viruses (Fig. 2A) (Dai et al., 2017; Pan *et al*., 2021). 5 major dsRNA segments are observed adjacent to each TEC and are labeled “Front Bottom,” “Bound,” “Front Top,” “Rear Top,” and “Rear Bottom” RNA segments based on their positions relative to the TEC (Fig. 2A). At the polar TECs a 6^th^ RNA molecule is observed adjacent to the TEC template entrance, labeled “Terminal” (Fig. 2B). Consistent with previous ARV structures, VP2 is a 1274-residue RNA-dependent RNA polymerase (RdRp) which, like CPV RdRp VP2, is organized into three major domains—N-terminal domain (1-387), C-terminal bracelet (902-1274), which sandwich the RdRp core (388-901), and are further differentiated into the thumb (793-901), fingers (388-557 and 595-690), and palm (558-594 and 691-792) subdomains (Fig. 2B-C) (Ding *et al*., 2018; Wang *et al*., 2018). The fingers house the NTP entry channel and, with the thumb, assist in elongation and proofreading while the palm catalyzes phosphodiester bond formation between new NTPs and the growing strand via D591, D740, and D741, all three of which appear in beta sheets and are highly conserved in reovirus RdRps (Fig. 2C) The N-and C-terminal domains are versatile: In addition to shielding the RdRp core from any internal environmental changes, they bind dsRNA and other viral core proteins (Fig. 3B,H). Consistent with our previous findings, the C-terminal bracelet of this non-transcribing ICP does not occlude the template exit channel as it does in quiescent CPV (Fig. 2G) (Ding *et al*., 2018). As previously observed, the polymerase possesses several channels to funnel RNA templates and transcription products (Movie S3) (Parker et al., 2002).

**Figure 2.**
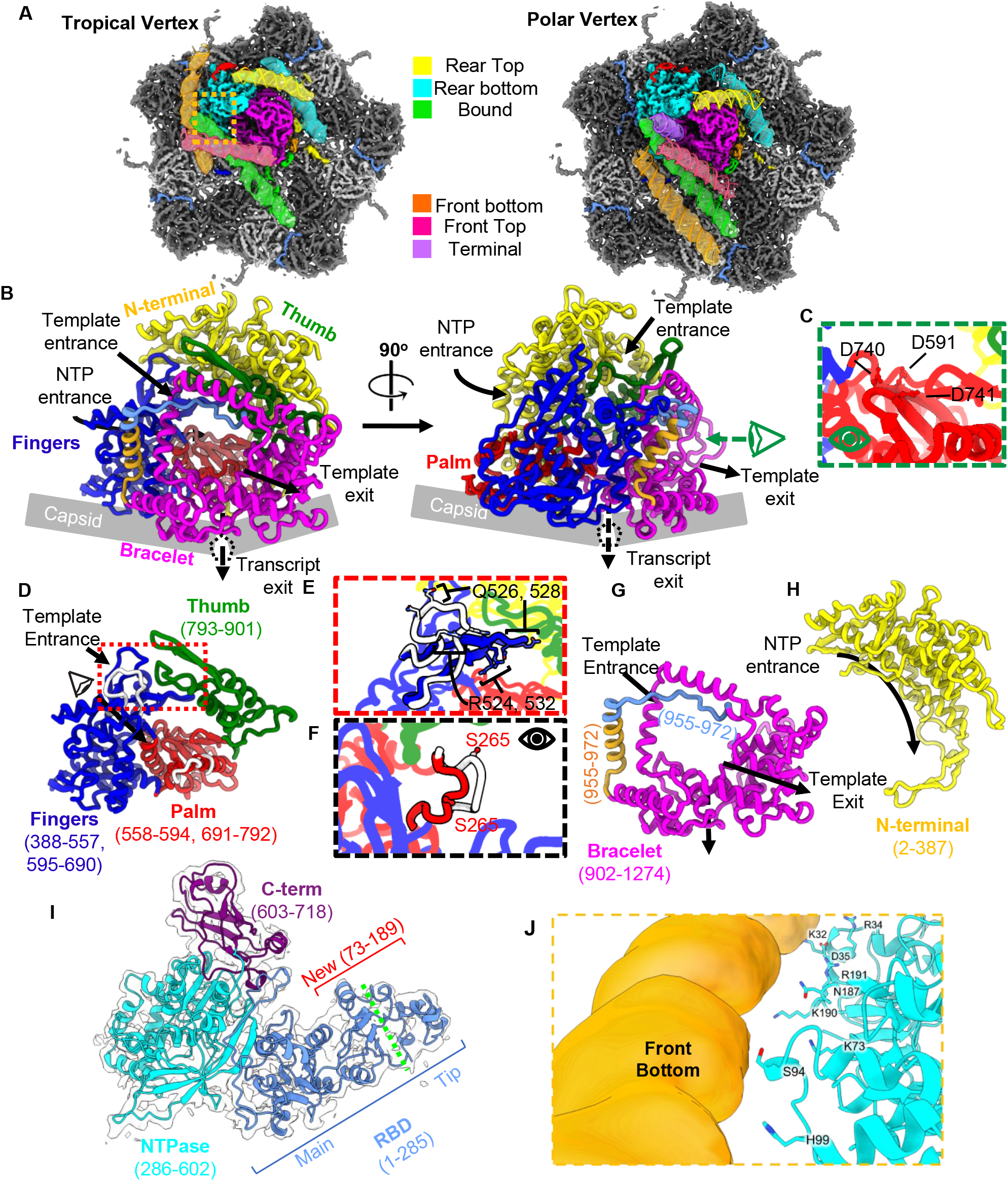
CryoEM maps show distinct RNA arrangement around different ICP vertices with identical TEC structures. (A) Comparison of the CryoEM density maps from polar and tropical vertices, highlighting the TEC (magenta and cyan) and surrounding RNA, shown semi-transparently and superposed with corresponding dsRNA models (colored by segment). (B-H) Atomic model of ICP RdRp VP2 colored by domain, either shown together (B), or separately (D-H) for detailed features. (C) provides magnified view of catalytic palm domain from B, with catalytically important residues indicated. In E and F views from D are shown to demonstrate the difference between the ICP and ISVP of the template channel finger loop (E) and catalytically important priming loop (F). (I) Atomic model of RdRp cofactor VP4 colored by domains with newly resolved residues indicated (red) and division of RBD into Main and Tip subdomains (green separator). (J) Interactions of the polar and charged residues from the newly resolved NTPase residues with the adjacent RNA segment from boxed region in A (orange).

**Figure 3.**
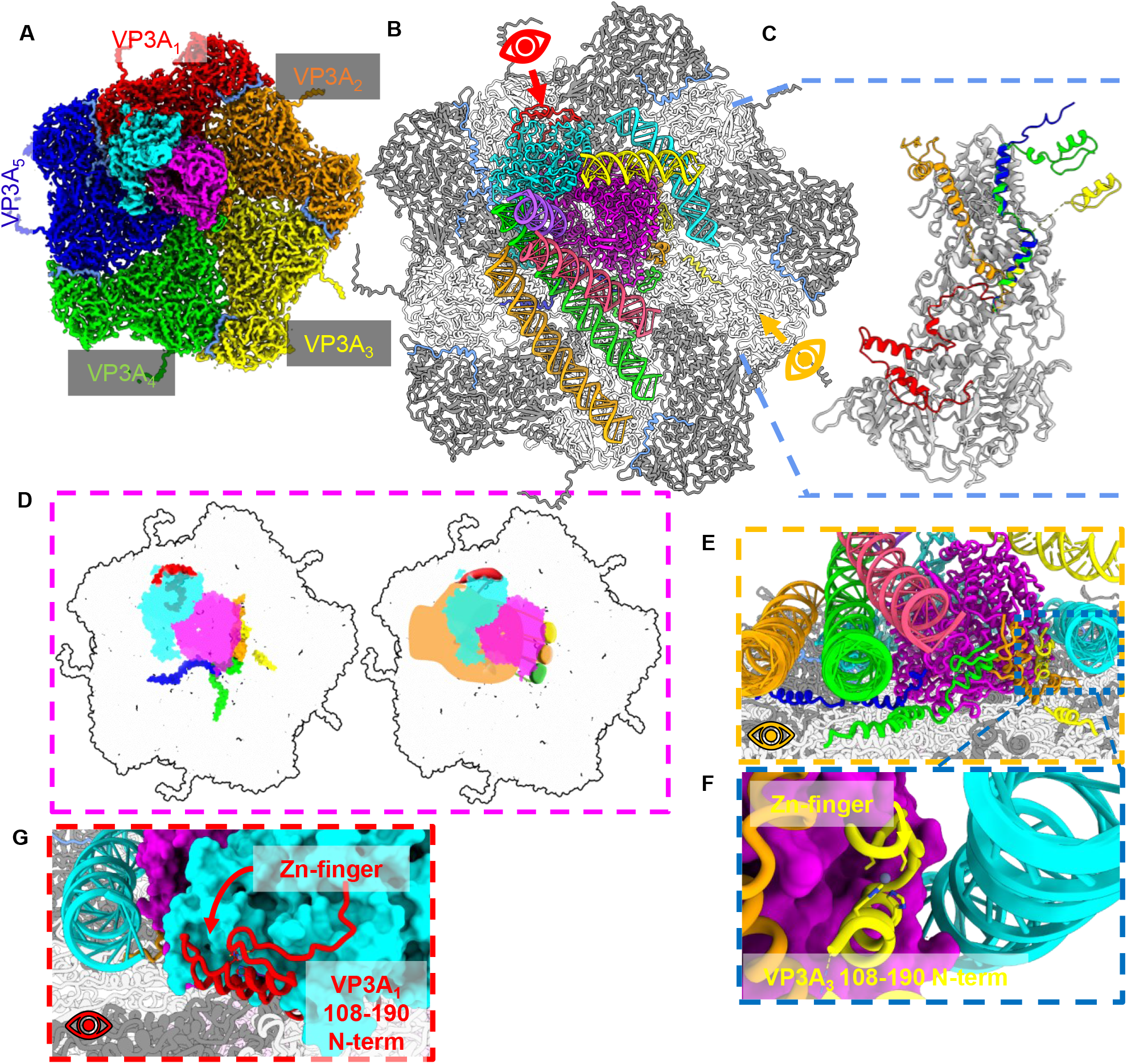
Asymmetric interactions of VP3_A_ N-termini with TEC and nucleic acids. (A and B) CryoEM map of polar decamer and attached TEC (left) with dimer pairs colored based on TEC association (A), and corresponding atomic model (B). Note the RNA is only shown in the atomic model (B) and not shown in the cryoEM density for clarity. (C) Superposition of VP3_A_ proteins from each dimer pair, with N-terminal residues colored to match dimer pairs from A. (D) Depiction of TEC along decamer lumen with N-terminal surfaces of VP3_A_ CSPs colored according to dimer pair in A (left), and hand shape added to improve visualization of Zn-fingers grasping TEC. (E-G) Side views of atomic model as indicated by eye symbols in B, with VP3_A_ N-termini colored as in C (E and G), and magnified view of VP3_A_3 Zn-finger domain interacting with both RdRp (magenta) and rear bottom dsRNA (cyan).

Our RdRp structure reveals several local architectural changes within these RdRp channels when compared to structures form the ARV ISVP (Fig 2D-E, and Movie S3). When viewed down the template entry channel we observe a positively charged Finger domain loop (a.a.s 520-535) moves further into the template entry channel from the ISVP to ICP, widening the channel for template entry (Fig. 2E, Movie S3). From within the channel, we observe the priming loop (a.a. 560-572), thought to separate template and transcript strands, moves away from the transcript exit channel and orients the catalytically important S267 residue towards the would-be incoming template (Fig. 2 F, Movie S3). This migration away from the bracelet domain also widens the mRNA exit channel, likely promoting exit through the adjacent I5 pores and turrets. From the external TEC view, we also observe an RdRp expansion along the capsid lumen appears linked to the radial expansion of the capsid beneath it (Movie S3). This contrasts with the VP4 NTPase which undergoes a unidirectional shift, consistent with the radial expansion of the capsid monomers with which it associated (Movie S3). These movements may be linked to the asymmetric association of the TEC along the expanding decameric subunit. As the NTPase is situated primarily atop the VP3_1_ dimer it moves along with the VP3_1_ dimer elongation, whereas the RdRp, which is seated across VP3_1-4_, is seemingly drawn in several directions based on the expansion of the capsid and the proportion of RdRp associated with each conformer pair (Fig. 3A). As uncoating is necessary to synthesize complete viral transcripts (Farsetta *et al*., 2000), the subtle conformational changes in the TECs observed here may be essential to carrying out efficient viral transcription.

### CSP VP3 and NTPase VP4 N-termini interact with replicative machinery

In the asymmetrically reconstructed sub-particles, the previously unmodeled RNA-binding domain in TEC NTPase are resolved alongside the Zn-finger containing N-terminal domains of VP3 CSPs each of which exhibit interactions with the viral genome (Fig. 3). We will first describe the newly resolved N-terminal region of the ARV NTPase.

Viral TECs are situated beneath 11 of the 12 I5 vertices in ARV, containing one RdRp and accompanying NTPase molecule per TEC (Fig. 1E). We previously reported the structure of the ARV NTPase, revealing three distinct domains: the N-terminal RNA-binding domain (RBD, a.a. 1-285), the NTPase domain (a.a. 286-602), and the C-terminal domain (CTD a.a. 603-718) (Ding *et al*., 2018). The NTPase RBD is composed of two subdomains, the ‘tip’ (a.a. 80-123) and ‘main’ (a.a. 1-79 and 124-285) which possess distinct electrostatic potentials (Fig. S3). In the previous structure, the *tip* domain was entirely unresolved along with a large portion of the *main* domain (a.a. 73-189) (Fig S3). In our structures we observe these previously missing NTPase segments extend away from the globular body of the TEC complex. This flexibility may help accommodate dynamic RNA interactions throughout transcription. The newly modeled N-terminal residues reveal a flexible region homologous to that of ARV’s MRV cousin but of a distinct fold (Pan *et al*., 2021), exhibiting limited RdRp contacts but extensive genome interactions (Fig 2I-J).

Previously, the N-terminal residues of VP3_A_ conformers were shown to associate with and lie along the exterior of the TEC, on both the RdRp and NTPase (Ding *et al*., 2018), and were suggested to anchor the TEC into place. The N-terminal region of ARV and MRV CSPs are also thought responsible for binding dsRNA during virus assembly (Lemay and Danis, 1994). The newly modeled VP3_A_ N-terminal residues include the previously unresolved residues 108-152 (VP3_A1-4_) which contain previously identified Zn-finger domains (a.a. 116-141) (Fig 3C-G), and conform to a traditional Cys_2_His_2_ motif, known to bind nucleotides (Fig 3E). Previous reporting suggests that the N-terminal Zn-fingers of MRV CSPs are unnecessary for dsRNA binding, and instead mediate stability of the capsid, despite demonstrated nucleotide binding affinity (Lemay and Danis, 1994; McDonald and Patton, 2011; Yu et al., 2011). In keeping with this we observe the four newly resolved Zn-fingers (VP3_A1-4_) lie along the TEC, with VP3_A2-4_ situated along an RdRp cleft opposite the NTPase, and VP3_A1_ seated along the VP4 NTPase domain forming a four-pronged setting that anchors the TEC complex to the capsid shell (Fig 3D). The VP3_A5_ N-terminus also extends towards the RdRp, however we were unable to resolve the Zinc finger in either sub-particle reconstruction. We also observe that at both the tropical and polar vertices the VP3_A3_ Zn-finger contacts the rear bottom genome segment (Fig 3E-F).

This observation suggests these N-terminal Zn-fingers may help to regulate transcription initiation as previously observed in rotavirus (Ding et al., 2019). It may be the case therefore that ARV VP3s are multifunctional, promoting TEC assembly and stability while maintaining genome organization in the quiescent particles. Despite the significant conformational changes of the capsid shell, the Zn-fingers appear in positions consistent with the low resolution ISVP structures, suggesting that the Zn-fingers may function independent of the capsid shell. This may be enabled by the flexible linker within the of the CSP N-terminal domains (a.a. 140-190), which extend slightly to accommodate the capsid expansion, while maintaining their TEC positioning and without altering the genome organization or activation state of the TEC (Movie S3).

### Structure of the ICP turret protein VP1, a multi-enzyme complex

Atop each I5 vertex of GSRV sits a pentameric turret composed of 5 copies of VP1 TPs (Fig 1A & 4A). These pentameric turrets form axial channels atop the pores through which the genome is translocated and provides mRNA capping functionality (Reinisch et al., 2000). CPV and MRV virions both have spike proteins which occupy the turret channel and serve dual roles, mediating cell entry, and occluding the mRNA exit channel to prevent premature transcript escape (Figure 4A) (Pan *et al*., 2021; Zhang *et al*., 2022b). ARVs lack a spike homologue therefore it is unclear how they might prevent premature transcript escape.

**Figure 4.**
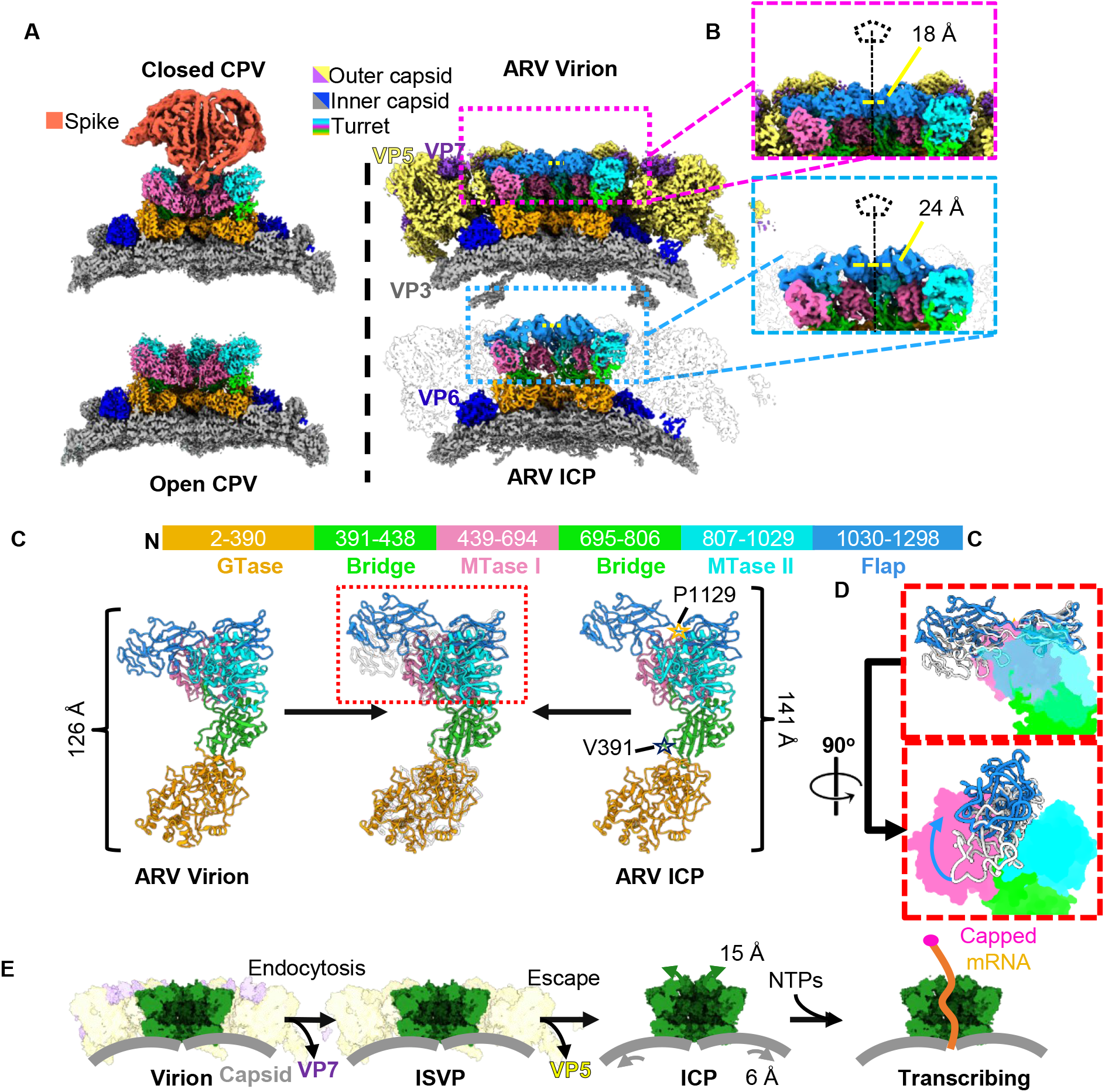
The novel conformational changes of the capping enzyme turret in uncoated virus. (A) comparison of CPV before and after detachment, where the Receptor binding spike remains attached to the turret and corresponding region of ARV virion and ICP. (B) Magnified view of turret pores from ARV virion (top) and ICP (bottom) with I5 symmetrical axis indicated with pentagon and line segment. (C) Atomic structure of the ARV turret protein VP1 from virion (left) and ICP (right) with superimposition of MTase domains (center) to show difference, with domains depicted (top) with the primary sequence number and corresponding color scheme. (D) magnified view of boxed region from C, with solid color of non-flap domains used to highlight virion to ICP differences. (E) Schematic demonstrating the confirmational changes differences undergone by ARV turrets (green) from virion to ISVP and ICP, which coincide with the loss of outer capsid proteins VP5 and VP7 (ghost yellow and purple).

Like MRV, ARV TPs are composed of five domains: the GTase (a.a. 2-390), a bridge domain (a.a. 391-438, 695-806), MTase I (439-694) MTase II (807-1029), and Flap (1030-1298) which contains 3 immunoglobulin-like (IG) folds, two of which extend to cover the axial channel (Fig 4C). When ICP and virion turrets are compared we observe a significant axial extension of the ARV turrets and widening of the turret channels (Fig. 4B-C). Local alignment of the 5 TP domains reveals that the enzymatic GTase and MTase domains remain stable (RMSD 0.842 Å and 0.907 respectively) while domain separation occurs at the flexible linker regions (a.a. 391-438, 695-806) between the GTase, MTase, and flap domains. This separation is enabled by the loss of the OCP VP5 that would otherwise clash with the new TP architecture (Fig 4A). Despite MRV possessing a similar double layered architecture, when the crystal structure of the MRV ICP is compared to the recently resolved asymmetric reconstruction of the complete virions the turret does not undergo analogous changes.

As ARV lacks a spike protein homologue, we investigated the flap domains that line the turret exit channel. When local alignment is performed it is clear there are significant conformational changes within the domain (Fig. 4 D). This difference is caused by the movement of the two C-terminal IG domains moving at the flexible linker region (a.a. 1129), which undergoes an outward shift of 14 Å and counterclockwise twist when viewed from within the I5 channel (Fig. 4D). This finding, coupled with the changes to capsid and TEC architecture, suggest a regulatory mechanism of transcription wherein the uncoating of ARV yields a particle whose capsid better accommodates genome transcription and whose capping enzymes are widened, or primed, for transcript export.

## Conclusions

This work provides the first near-atomic resolution asymmetric reconstruction of an ICP from a turreted, multilayered reovirus particle which exhibits subtle but functionally important conformational changes compared to coated ARV structures. These changes in the internal capsid volume, along-side a widening of TEC nucleotide channels and extension of the 5’ -cap synthesizing turret proteins create the architectural environment conducive to endogenous transcription. As the ICP genome remains in the quiescent state, it is conceivable that transcription initiation requires factors such as S-adenosyl methionine and NTPs in addition the conformational changes observed here. In this manner, the highly specialized outer capsid layers serve as a useful transcriptional regulator, ensuring that transcription requires not only the presence of cofactors, but also loss of outer capsid layers which occurs upon cell entry.

## Acknowledgements

This work was supported in part by grants from the National Institutes of Health (AI094386 to Z.H.Z.). A.S. received support from NIH Ruth L. Kirschtein National Research Service Award AI007323. We acknowledge the use of resource at the Electron Imaging Center for Nanomachines supported by UCLA and by instrumentation grants from NIH (1S10RR23057, 1S10OD018111 and U24GM116792) and NSF (DBI-1338135 and DMR-1548924). We acknowledge support from the UCLA AIDS Institute, the James B. Pendleton Charitable Trust and the McCarthy Family Foundation.

## Author contributions

Z.H.Z. designed and supervised the project; S.S. prepared sample and made cryoEM grids; Y.C. performed cryoEM imaging and 3D reconstruction; A.S. built the atomic models, interpreted the structures, made the figures, and wrote the paper with input from Z.H.Z.; All authors reviewed and approved the paper.

## Conflict of interest

The authors declare no competing interests.

## Methods and Data Availability

### Viral Culture and Isolation

Golden shiner reovirus stock was purchased from ATCC (ATCC® VR957™) and was propagated in Fat Head Minnow cells (FHM; ATCC® CCL-42™) with a slight modification to previously described method (Shaw *et al*., 1996). FHM cells were cultured in MEM, hanks’ balanced salts (Gibco™) supplemented with 10% fetal bovine serum (Atlanta biologicals), Penicillin (10,000 IU) and Streptomycin (10,000 µg/ml) (Corning ®, USA), and maintained at 28°C. Confluent monolayers of FHM were infected with ∼5 plaque forming units of GSRV per cell for ∼48 hours. Once cell syncytia was visible, media was collected from the infected cell culture and centrifuged at 11,000 g for 15 min to get rid of cells and cellular debris. Clean supernatant was centrifuged at 100,000g for 3 hours (SW28 rotors, Beckman Coulter) to pellet the viral particles. Supernatant was discarded and 100 μl of ice-chilled PBS buffer was added to each pellet to resuspend the viral particles overnight. Pellets were pooled and loaded on a 15%-50% sucrose gradient and banded by centrifuging at 100,000 g for 1 hour (SW41 rotor) at 4°C. Band containing the GCRV cores was harvested and diluted in PBS, pelleted at 100,000 g for 2 hours. The cores in the pellet were resuspended in PBS and quality confirmed by negative-staining (2% uranyl acetate) stain transmission electron microscopy.

### CryoEM imaging

For cryoEM sample preparation, 2.5-μL aliquots of the GSRV core sample were applied to 200-mesh Quantifoil grid with 2µm-diameter hole size (2/2 or 2/1 Quantifoil) and plunge frozen in a Vitrobot Mark IV (Thermo Fisher Scientific) (Thermo Fisher Scientific) after blotting (blot time 15 seconds, blot force -5, 100% humidity and 4°C). Movies of dose fractionated image frames were recorded using a Titan Krios microscope (Thermo Fisher Scientific) equipped with a Gatan imaging filter (GIF) quantum LS and a post-GIF Gatan K2 Summit direct electron detector operated in super resolution mode with a calibrated pixel size of 0.68 Å/pixel with SerialEM (Mastronarde, 2005). The GIF slit width was set to 20 eV at the zero-loss peak, and dose rate as recorded by the camera was set to with a total exposure dosage of ∼45 e/Å^2^ at the sample level which was fractionated equally into 40 frames. A defocus range of 1.5µm-2.5 µm was targeted for recordings. Dose-fractionated frames were binned 2× (effective pixel size 1.36 Å) and aligned for beam induced motion correction to produce dose-weighted and unweighted micrographs, used for final reconstruction and initial screening and CTF estimation respectively, using MotionCor2 (Zheng et al., 2017). In total 12,258 movies were recorded.

### Data processing

Defocus values of the micrographs were initially estimated using CTFFIN4 (Rohou and Grigorieff, 2015), and those micrographs with significant ice contamination or defocus values beyond -0.8 and -3.0 µm were removed leaving 10,935 micrographs. 38,486 particles were initially selected by Ethan (Kivioja et al., 2000). The particles were extracted and initially subjected to an icosahedral refinement in RELION which yielded a 3.4 Å cryoEM map of the complete viral particle (Scheres, 2012). In our icosahedral reconstruction, strong densities were observable beneath the positions of I5 vertices. We then followed a previously established stepwise symmetry relaxation workflow within RELION to generate asymmetric sub-particle reconstructions of both polar and tropical vertices (Ding *et al*., 2018; Scheres, 2012; Zhang *et al*., 2015). Briefly, particles from the 3.2 Å icosahedral reconstruction were subjected to symmetry expansion (option I3) using the *relion_symmetry_expand* command, to generate a RELION STAR file containing 60 orientation entries for each ARV particle, corresponding to 5 copies of each I5 vertex differentiated by the rotation about the I5 vertex, and listed as _rlnAngleRot. One entry corresponding to each vertex was kept, yielding a duplicate-entrues-removed STAR file with ∼12 entries, or one for each vertex, for each ICP. The icosahedral reconstruction orients in such a way that the location of one vertex consistently lies along the *z* axis, so we performed classification on ICP particles with the capsid density subtracted to identify tropical and polar vertices and estimated their location for extraction of the penton sub-particles using the RELION “Particle extraction” tool (Scheres, 2013). Polar and Tropical vertices were separately refined into the final sub-particle maps provided here. Unfortunately, this method failed to resolve the true asymmetric structure featuring 11 TECs, as the pseudo-D3 symmetrical particle featured 12 TECs, hindering efforts to resolve the unoccupied vertex. This was overcome following the D3-symmetry expansion of the ICP particles, generating 6 duplicates within a new STAR file. The particles were then subjected to exhaustive classification with a reference mask beneath a tropical vertex to identify a class without a TEC. Using the orientations determined during this classification we were able to reconstruct the complete asymmetric ICP.

### Atomic Model Building and Refinement

Atomic models of the GSRV RdRp, CSP, NTPase, turret and clamp proteins were initially modeled into the cryoEM density in Coot using the GCRV ISVP (PDB 6M99) as a homology following previously established protocols (Ding *et al*., 2018; Emsley et al., 2010; Yu et al., 2018). Following initial fitting, the polypeptide chains were mutated to match the GCRV sequence using Coot’s Mutate Residue Range tool prior to local and global refinement in PHENIX (Afonine et al., 2018). Residues were built into previously unresolved residues, de novo using Coot, except for the first 200 residues of VP4, whose structure was predicted in AlphaFold2 prior to fitting into the cryoEM map (Jumper et al., 2021). Upon initial fitting the model was refined within the sub-particle map using the molecular dynamic flexible fitting (MDFF) feature in ISOLDE from the UCSF ChimeraX user interface (Croll, 2018; Goddard *et al*., 2018). Models were then subjected to a final round of PHENIX real space refine and the resulting structure was validated by the worldwide protein data bank validation server (Afonine *et al*., 2018; Berman et al., 2003). Root mean squared deviations (RMSDs) between protein chains were calculated using the matchmaker tool in UCSF ChimeraX. Internal capsid volume was calculated using the ChimeraX recipe “Measure volume enclosed by a virus capsid” on our ARV ICP and the previously published virion structure (EMD-6969) (Wang *et al*., 2018). TEC distribution maps were generated using UCSF Chimera (Pettersen et al., 2004).

### Modeling of dsRNA

We inserted poly-AU, A-form dsRNA segments from a previously published MRV model (PDB 7ELH) into our cryoEM maps (Pan *et al*., 2021). Segments were roughly placed into the cryoEM map, and segments were further fit in ISOLDE (Croll, 2018). Briefly, adaptive distance restraints were applied to the nucleic acid segments with reduced weight applied to the non-crystallographic map were fit into the cryoEM maps using ISOLDEs MDFF feature (Croll, 2018).

### Data Availability

CryoEM maps of the asymmetric GSRV ICP and sub-particles have been submitted to the Electron Microscopy Data Bank and can be found under accession numbers EMD-XXXXX, EMD-XXXXX and EMD-XXXXX. The coordinates of ICP sub-particle were deposited in the Protein Data Bank under accession number XXXX.

## Figure Legends

**Figure S1.**
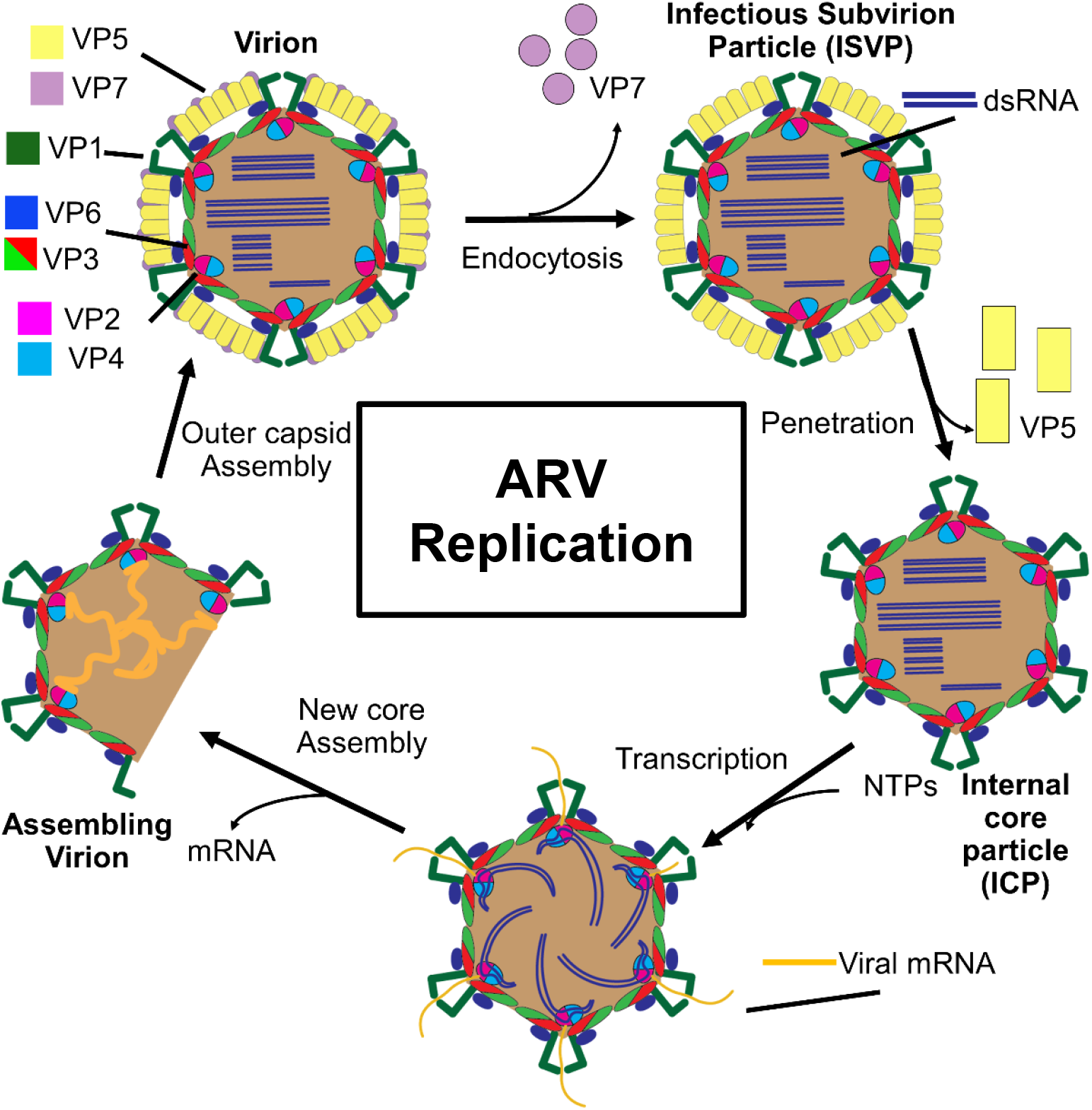
Typical changes to Aquareovirus capsid during replication. Diagrammatic representation of an Aquareovirus in cross section and the proposed replication process. The identities of viral structural and enzymatic proteins are indicated using the nomenclature for Aquareoviruses and assigned colors. Double-stranded DNA is represented by parallel blue line segments and single-stranded RNA is represented by orange lines.

**Figure S2.**
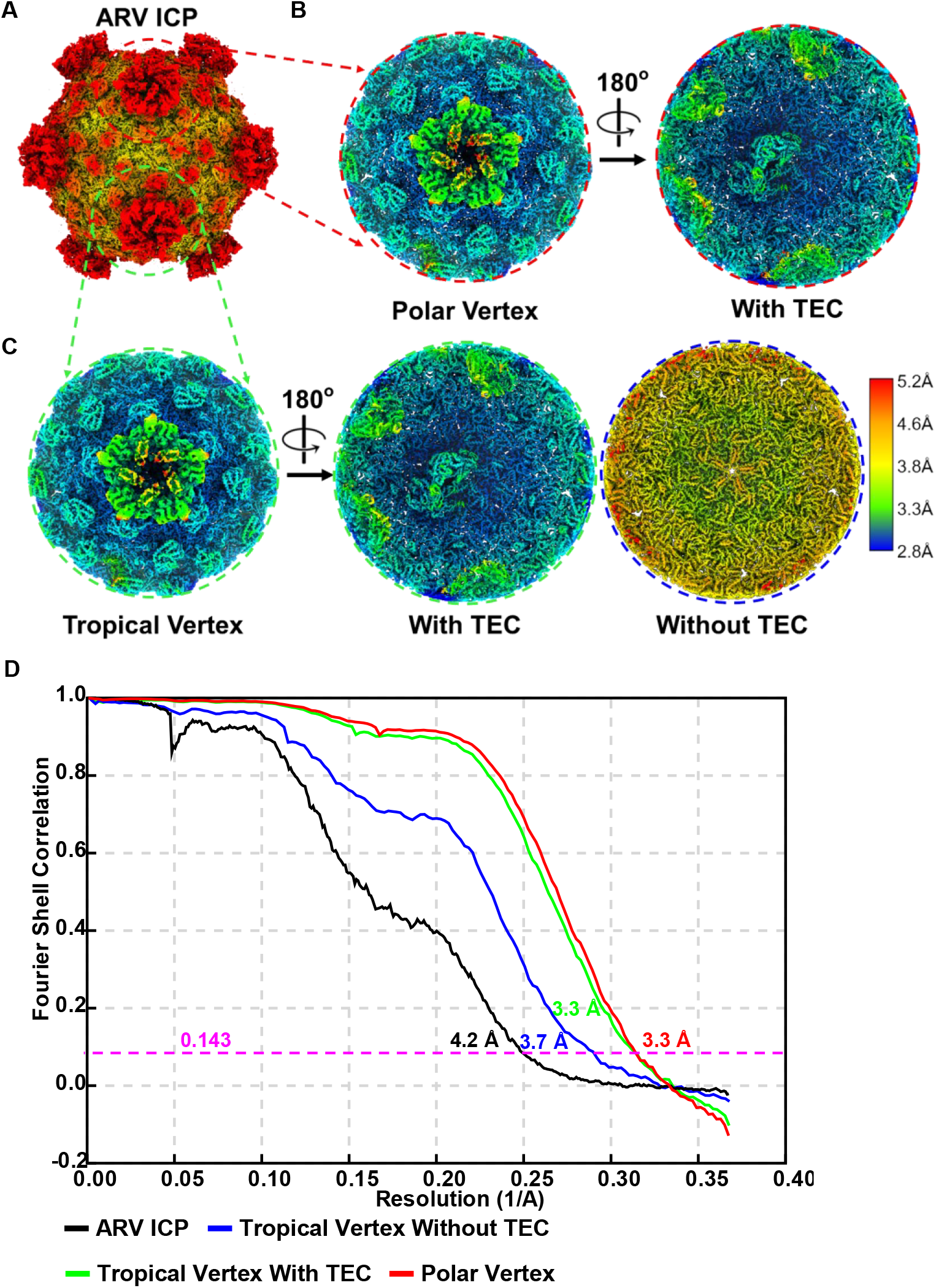
Asymmetric reconstructions from ARV ICP. (A-C) CryoEM maps of GSRV ICP (A) and sub particles from Polar (B) and Tropical (C) vertices colored by local resolution. Internal views of Polar and Tropical vertices in B and C (right and middle panels respectively) showing the asymmetrically associated TEC in both species and the lack thereof in a subset of Tropical vertices (C, right). (D) Gold-standard Fourier shell correlation (FSC) curves calculated from the independently refined half-maps of each cryoEM reconstruction in A-C, with resolution values listed for each species at FSC = 0.143.

**Figure S3.**
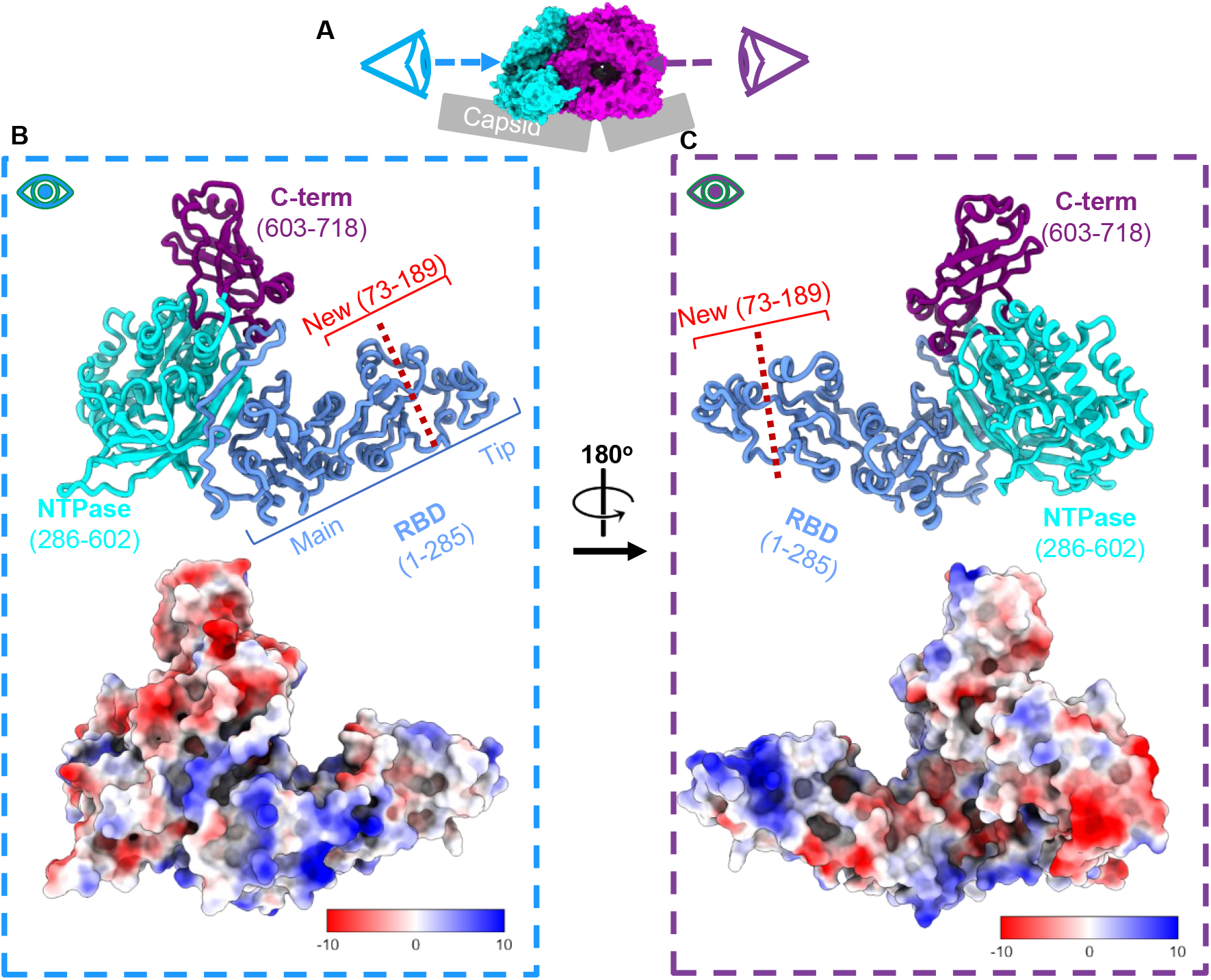
Columbic potential maps of NTPase VP4. (A-C) different views of NTPase VP4 protein depicted by ribbon diagrams and colored by domains (B-C, top). (B) View of NTPase as viewed along the face opposite the associated RdRp with surface colored by columbic potential (bottom). (C) View of the RdRp associated face of NTPase depicted in the same manner as B.

**Movie S1.**
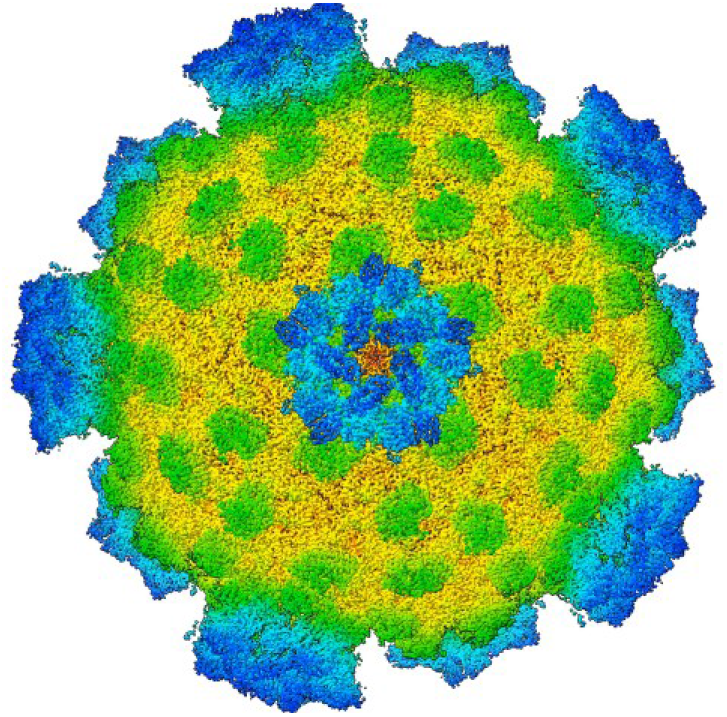
Expansion of ARV ICP decamer. Comparison of cryoEM maps and atomic models from core of ARV virion particle and ICP. Cartoon representation of ARV ICP, without TEC, conforming to same color scheme used in Fig. 1A. First morph movie demonstrates rise away from capsid center. Second morph toggles between virion and ICP decamers, superimposed at residues adjacent to the I5 vertices, from internal, side and external view to highlight conformational changes of capsid shell proteins (lime and red) and capping enzyme proteins (dark green). Rising motion away from virion origin not shown.

**Movie S2.**
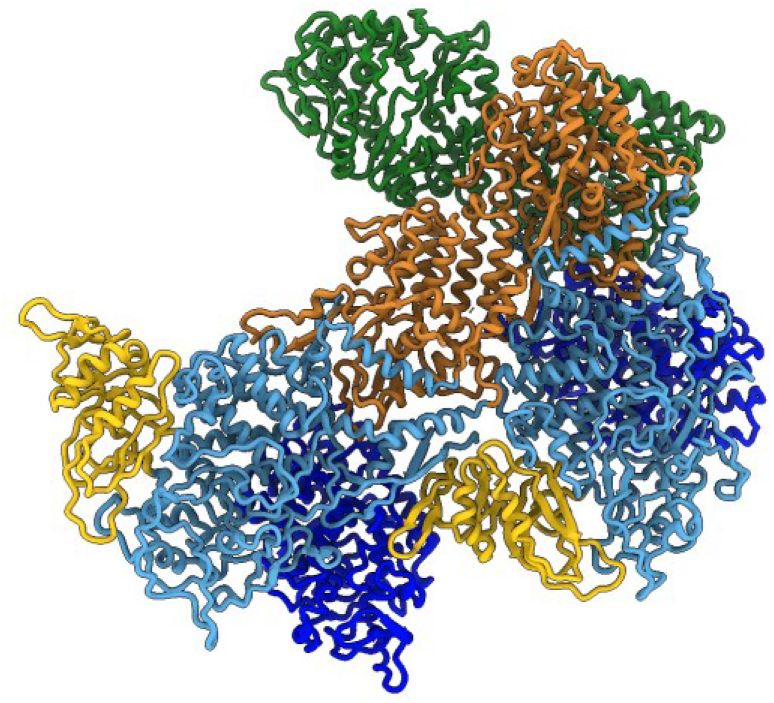
Local motion of CSP dimer from Virion to ICP. Cartoon representation of CSP dimer as colored in Fig. 1D, with local VP6 clamp proteins and the first 390 residues of the Turret protein VP1. Demonstrates local motion of VP3 monomers morphing between virion and ICP conformations. Side view demonstrates radial extension of VP3_A_ and VP3_B_ with the conserved position of VP3_A_ a-helices 13 and 14. External view shows radial movement of clamp and turret proteins along external wall of ICP while their removal demonstrates the constrained movement of their interfaces during the morph.

**Movie S3.**
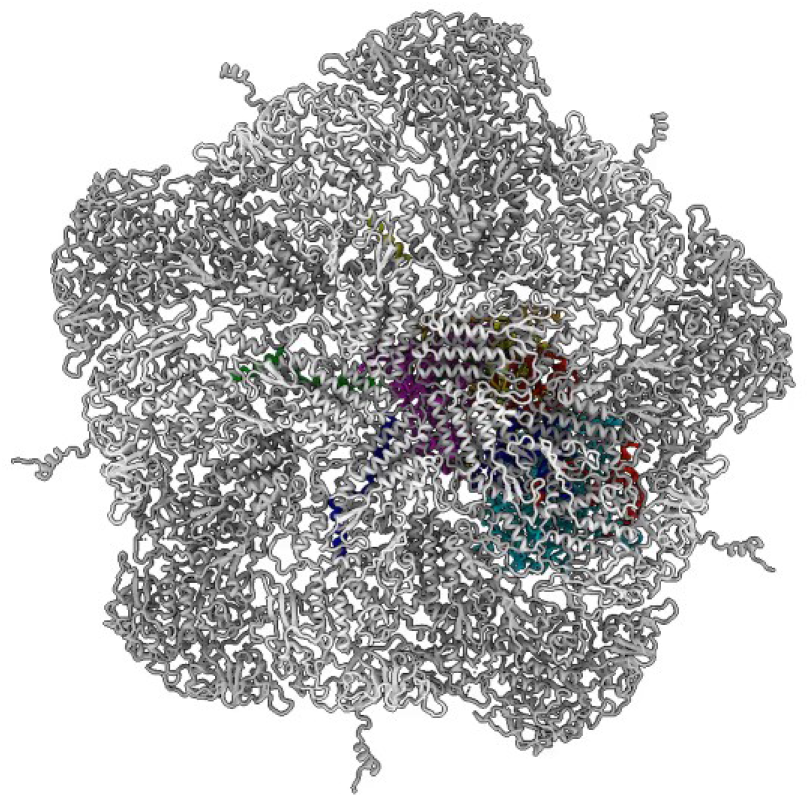
Conformational changes from ISVP to ICP TEC. Cartoon representation of decamer and TEC morphs between ISVP and ICP, demonstrating asymmetric movement of TEC, with NTPase moving along with capsid lumen while the RdRp appears to expand, creating a breathing like motion. Movie also emphasizes changes to RdRp finger loop at template entry channel and priming loop retraction from the transcript exit channel.

**Movie S4.**
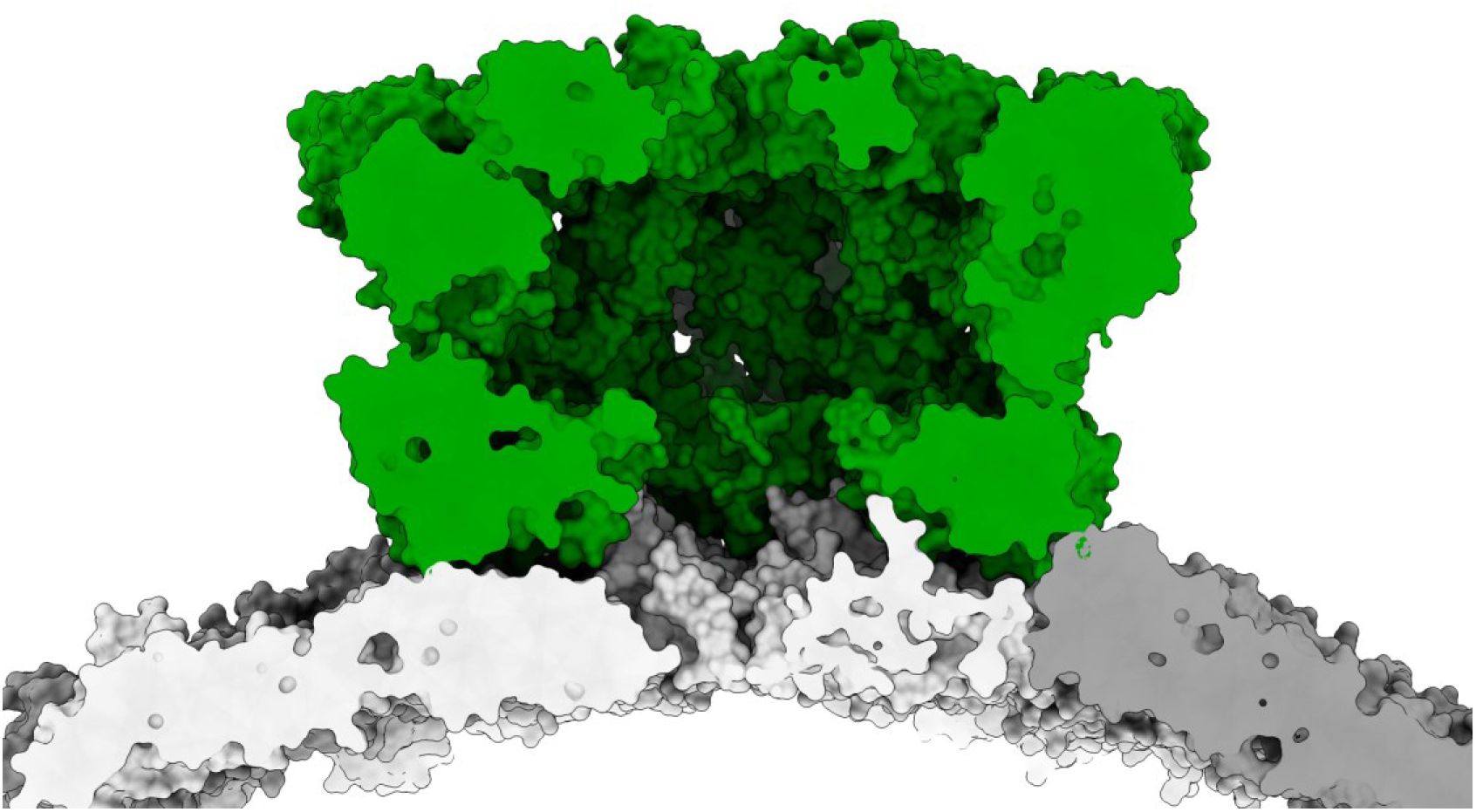
Extension of Turret proteins. Cross-sectional view of ARV turret proteins morphing between virion and ICP structures along fixed CSPs from the ICP. Movie begins with virion turret proteins represented as surfaces morphing to ICP conformations.

## Notes

### Competing Interest Statement

The authors have declared no competing interest.

### Summary of Updates

Revision has been made to improve clarity

## References

Afonine, P.V., Poon, B.K., Read, R.J., Sobolev, O.V., Terwilliger, T.C., Urzhumtsev, A., and Adams, P.D. (2018). Real-space refinement in PHENIX for cryo-EM and crystallography. Acta Crystallographica Section D: Structural Biology. International Union of Crystallography.

Berman, H., Henrick, K., and Nakamura, H. (2003). Announcing the worldwide Protein Data Bank. Nature Structural & Molecular Biology.

Chen, F., Zhu, L., Zhang, Y., Kumar, D., Cao, G., Hu, X., Liang, Z., Kuang, S., Xue, R., and Gong, C. (2018). Clathrin-mediated endocytosis is a candidate entry sorting mechanism for Bombyx mori cypovirus. Scientific Reports.

Croll, T.I. (2018). ISOLDE: A physically realistic environment for model building into low-resolution electron-density maps. Acta Crystallographica Section D: Structural Biology. International Union of Crystallography.

Cui, Y., Zhang, Y., Zhou, K., Sun, J., and Zhou, Z.H. (2019). Conservative transcription in three steps visualized in a double-stranded RNA virus. Nature Structural & Molecular Biology. Springer US.

Dai, X., Li, Z., Lai, M., Shu, S., Du, Y., Zhou, Z.H., and Sun, R. (2017). In situ structures of the genome and genome-delivery apparatus in a single-stranded RNA virus. Nature. Nature Publishing Group.

Demidenko, A.A., and Nibert, M.L. (2009). Probing the transcription mechanisms of reovirus cores with molecules that alter RNA duplex stability. Journal of virology.

Ding, K., Celma, C.C., Zhang, X., Chang, T., Shen, W., Atanasov, I., Roy, P., and Zhou, Z.H. (2019). In situ structures of rotavirus polymerase in action and mechanism of mRNA transcription and release. Nature Communications. Springer US.

Ding, K., Nguyen, L., and Zhou, Z.H. (2018). In Situ Structures of the Polymerase Complex and RNA Genome Show How Aquareovirus Transcription Machineries Respond to Uncoating. In S. López, ed. Journal of Virology.

Dryden, K., Wang, G., Yeager, M., Nibert, M., Coombs, K., Furlong, D., Fields, B., and Baker, T. (1993). Early steps in reovirus infection are associated with dramatic changes in supramolecular structure and protein conformation: analysis of virions and subviral particles by cryoelectron microscopy and image reconstruction. Journal of Cell Biology.

Ebert, D.H., Deussing, J., Peters, C., and Dermody, T.S. (2002). Cathepsin L and Cathepsin B Mediate Reovirus Disassembly in Murine Fibroblast Cells. Journal of Biological Chemistry. 2002 ASBMB. Currently published by Elsevier Inc; originally published by American Society for Biochemistry and Molecular Biology.

Emsley, P., Lohkamp, B., Scott, W.G., and Cowtan, K. (2010). Features and development of Coot. Acta Crystallographica Section D Biological Crystallography. International Union of Crystallography.

Fang, Q., Shah, S., Liang, Y., and Zhou, Z.H. (2005). 3D reconstruction and capsid protein characterization of grass carp reovirus. Science in China, Series C: Life Sciences.

Farsetta, D.L., Chandran, K., and Nibert, M.L. (2000). Transcriptional Activities of Reovirus RNA Polymerase in Recoated Cores. Journal of Biological Chemistry. 2000 ASBMB. Currently published by Elsevier Inc; originally published by American Society for Biochemistry and Molecular Biology.

Goddard, T.D., Huang, C.C., Meng, E.C., Pettersen, E.F., Couch, G.S., Morris, J.H., and Ferrin, T.E. (2018). UCSF ChimeraX: Meeting modern challenges in visualization and analysis. Protein Science.

Hill, C.L., Booth, T.F., Prasad, B.V.V., Grimes, J.M., Mertens, P.P.C., Sutton, G.C., and Stuart, D.I. (1999). The structure of a cypovirus and the functional organization of dsRNA viruses. Nature Structural Biology.

Jumper, J., Evans, R., Pritzel, A., Green, T., Figurnov, M., Ronneberger, O., Tunyasuvunakool, K., Bates, R., Žídek, A., Potapenko, A., et al. (2021). Highly accurate protein structure prediction with AlphaFold. Nature. Springer US.

Kim, J., Tao, Y., Reinisch, K.M., Harrison, S.C., and Nibert, M.L. (2004). Orthoreovirus and Aquareovirus core proteins: conserved enzymatic surfaces, but not protein–protein interfaces. Virus Research.

King, A.M.Q., Adams, M.J., Carstens, E.B., and Lefkowitz, E.J. (2012). Family - Reoviridae. In A.M.Q. King, M.J. Adams, E.B. Carstens, and E.J. Lefkowitz, eds. Virus Taxonomy. Elsevier.

Kivioja, T., Ravantti, J., Verkhovsky, A., Ukkonen, E., and Bamford, D. (2000). Local average intensity-based method for identifying spherical particles in electron micrographs. Journal of structural biology.

Lemay, G., and Danis, C. (1994). Reovirus 1 Protein: Affinity for Double-stranded Nucleic Acids by a Small Amino-terminal Region of the Protein Independent From the Zinc Finger Motif. Journal of General Virology.

Mastronarde, D.N. (2005). Automated electron microscope tomography using robust prediction of specimen movements. Journal of Structural Biology.

McDonald, S.M., and Patton, J.T. (2011). Rotavirus VP2 Core Shell Regions Critical for Viral Polymerase Activation. Journal of Virology.

McEntire, M.E., Iwanowicz, L.R., and Goodwin, A.E. (2003). Molecular, physical, and clinical evidence that golden shiner virus and grass carp reovirus are variants of the same virus. Journal of Aquatic Animal Health.

Nason, E.L., Samal, S.K., and Venkataram Prasad, B.V. (2000). Trypsin-Induced Structural Transformation in Aquareovirus. Journal of Virology.

Pan, M., Alvarez-Cabrera, A.L., Kang, J.S., Wang, L., Fan, C., and Zhou, Z.H. (2021). Asymmetric reconstruction of mammalian reovirus reveals interactions among RNA, transcriptional factor µ2 and capsid proteins. Nature Communications. Springer US.

Pettersen, E.F., Goddard, T.D., Huang, C.C., Couch, G.S., Greenblatt, D.M., Meng, E.C., and Ferrin, T.E. (2004). UCSF Chimera-A visualization system for exploratory research and analysis. Journal of Computational Chemistry.

Reinisch, K.M., Nibert, M.L., and Harrison, S.C. (2000). Structure of the reovirus core at 3.6 Å resolution. Nature.

Rohou, A., and Grigorieff, N. (2015). CTFFIND4: Fast and accurate defocus estimation from electron micrographs. Journal of Structural Biology. Elsevier Inc.

Scheres, S.H.W. (2012). RELION: Implementation of a Bayesian approach to cryo-EM structure determination. Journal of Structural Biology. Elsevier Inc.

Scheres, S.H.W. (2013). Single-particle processing in RELION. Manuals.

Shaw, A.L., Samal, S.K., Subramanian, K., and Prasad, B.V. (1996). The structure of aquareovirus shows how the different geometries of the two layers of the capsid are reconciled to provide symmetrical interactions and stabilization. Structure.

Skehel, J.J., and Joklik, W.K. (1969). Studies on the in vitro transcription of reovirus RNA catalyzed by reovirus cores. Virology.

Trask, S.D., McDonald, S.M., and Patton, J.T. (2012). Structural insights into the coupling of virion assembly and rotavirus replication. Nature Reviews Microbiology.

Troeger, C., Khalil, I.A., Rao, P.C., Cao, S., Blacker, B.F., Ahmed, T., Armah, G., Bines, J.E., Brewer, T.G., Colombara, D.V., et al. (2018). Rotavirus Vaccination and the Global Burden of Rotavirus Diarrhea Among Children Younger Than 5 Years. JAMA pediatrics.

Wang, X., Zhang, F., Su, R., Li, X., Chen, W., Chen, Q., Yang, T., Wang, J., Liu, H., Fang, Q., and Cheng, L. (2018). Structure of RNA polymerase complex and genome within a dsRNA virus provides insights into the mechanisms of transcription and assembly. Proceedings of the National Academy of Sciences of the United States of America.

Yang, C., Ji, G., Liu, H., Zhang, K., Liu, G., Sun, F., Zhu, P., and Cheng, L. (2012). Cryo-EM structure of a transcribing cypovirus. Proceedings of the National Academy of Sciences of the United States of America.

Yu, I., Nguyen, L., Avaylon, J., Wang, K., Lai, M., and Zhou, Z.H. (2018). Building atomic models based on near atomic resolution cryoEM maps with existing tools. Journal of Structural Biology. Elsevier.

Yu, X., Ge, P., Jiang, J., Atanasov, I., and Zhou, Z.H. (2011). Atomic Model of CPV Reveals the Mechanism Used by This Single-Shelled Virus to Economically Carry Out Functions Conserved in Multishelled Reoviruses. Structure. Elsevier Ltd.

Yu, X., Jiang, J., Sun, J., and Zhou, Z.H. (2015). A putative ATPase mediates RNA transcription and capping in a dsRNA virus. eLife.

Zhang, F., Sun, D., and Fang, Q. (2022a). Molecular Characterization of Outer Capsid Proteins VP5 and VP7 of Grass Carp Reovirus. Viruses.

Zhang, H., Zhang, J., Yu, X., Lu, X., Zhang, Q., Jakana, J., Chen, D.H., Zhang, X., and Zhou, Z.H. (1999). Visualization of Protein-RNA Interactions in Cytoplasmic Polyhedrosis Virus. Journal of Virology.

Zhang, X., Ding, K., Yu, X., Chang, W., Sun, J., Zhou, Z.H., and Hong Zhou, Z. (2015). In situ structures of the segmented genome and RNA polymerase complex inside a dsRNA virus. Nature. Nature Publishing Group.

Zhang, X., Jin, L., Fang, Q., Hui, W.H., and Zhou, Z.H. (2010). 3.3 Å Cryo-EM Structure of a Nonenveloped Virus Reveals a Priming Mechanism for Cell Entry. Cell.

Zhang, Y., Cui, Y., Sun, J., and Zhou, Z.H. (2022b). Multiple conformations of trimeric spikes visualized on a non-enveloped virus. Nature Communications. Springer US.

Zheng, S.Q., Palovcak, E., Armache, J.-P., Verba, K.A., Cheng, Y., and Agard, D.A. (2017). MotionCor2: anisotropic correction of beam-induced motion for improved cryo-electron microscopy. Nature Methods.

Zhou, Z.H., Zhang, H., Jakana, J., Lu, X.-Y., and Zhang, J.-Q. (2003). Cytoplasmic Polyhedrosis Virus Structure at 8 A by Electron Cryomicroscopy: Structural Basis of Capsid Stability and mRNA Processing Regulation. Structure.

